# Quantifying structural vulnerabilities and resilience to research integrity risks in biomedical research networks

**DOI:** 10.64898/2026.06.26.734852

**Authors:** Zhongchao Pan, B. Ian Hutchins

## Abstract

Retractions of scientific articles reflect a rising research integrity risk in biomedical research, and continued citation of retracted articles poses a downstream risk to research that built upon unreliable studies. Few studies have examined the structural risk factors embedded in how the research enterprise is organized. Here, we apply an analytical risk assessment framework to structural risk factors associated with retracted articles and the propagation of their citations to ten broad fields of biomedical research, collectively accounting for 14% of biomedicine. We pronounced field-specific differences in retraction rates and citations to retracted articles, and show that multiple retractions from single authors occur far more often than chance would predict. By subdividing fields into finer-grained topics with machine learning network clustering, we find that individual authors can reach large proportions of the literature within their topics through citations, and that retraction risk is positively associated with author productivity. Across these fields, 22% of the literature cites authors with at least one retracted article, and 6% cite work from authors with multiple prior retractions. Together, these factors lead to a concentration of risk within topics, which is only partially explained by the uneven distribution of authors with multiple retractions. Despite these quantifiable vulnerabilities, the risk has not yet been fully realized: most topics cite retracted work no more than baseline. Our findings thus reveal both a latent network vulnerability to the rapid dissemination of questionable results and a measurable resilience that has so far kept this possibility in check. The scientific community would particularly benefit from targeted efforts to test and strengthen reproducibility in high-risk scientific topics.

## Introduction

Retractions remain rare events despite exponential growth in recent decades (1). Yet the research integrity risks associated with retractions attract substantially more attention than their frequency would imply (2). An example of this was the recent retraction of an Alzheimer’s Disease paper due to image manipulation (3). This called into question the validity of the Alzheimer’s field as a whole because that paper had been cited over 2000 times (4). Similar concerns have been raised about for retracted clinical papers, which can continue to be used and cited for a decade or more after their retraction (5,6). Research integrity risks attract disproportionate attention not because they are frequent, but because they call into question the validity of a potentially much larger body of work that built upon retracted papers (7–9).

The scientific community has tended to treat retractions as an individual failing. Misconduct is the most frequently cited reason why a paper is retracted (10,11). Moreover, multiple retractions are often attributed to misconduct from individual scientists (12). However, viewing research integrity risk as a structural issue may be more productive (13). Scientists conduct their work in a competitive enterprise, with norms regarding the conduct of research and applicable methodologies that vary from field to field. Incentives due to hypercompetition (14,15) and opportunities for misconduct may differ greatly depending on field (16). Furthermore, the extent of building upon prior work, and the attendant citation, may also differ by scientific area.

This means that the degree of vulnerability is likely to be a structural property of the system, and one that is measurable through the publication and authorship knowledge networks. For example, (17) measured the extent to which individual authors are cited in the field of Alzheimer’s Disease and found that top authors reach about 7.5% of that field through citations. In practice, the paper retracted due to research misconduct could have affected at most 2% of the field, suggesting that such risk is both readily measured and also that this latent risk has not been fully realized even in the most extreme examples. However, this kind of risk assessment has not been performed for other fields.

Because the magnitudes of structural research integrity risks are not known, in this study, we quantify the structural vulnerabilities and potential resilience to research integrity risks in research networks across a broad span of biomedical fields. By examining retraction rates across ten broad fields of biomedical research, we observe substantial field-level variance in the rate of retraction, and the rate at which retracted papers are cited within their respective fields. We show a substantially elevated risk of multiple retractions from the same author, and demonstrate a positive relationship between author productivity and odds of retraction. We show that overall retraction rates are a low proportion of the published literature in each field, but within fields, retractions tend to be concentrated in specific topics (identified by graph clustering of the citation network in each field). Finally, we observe substantial concentration of risk in research topics that is independent of the risk of multiple retractions, suggesting structural effects on research integrity risk at the subfield level.

## Results

Although retracted papers present the most visible form of research integrity problems in science, not all retractions are issued for reasons that call into question the validity of the results presented in the paper. Most retracted papers are thought to result from misconduct, such as data fabrication or image manipulation. However, papers may be retracted for other reasons, such as failure to comply with ethical review requirements, or because of issues related to credit, such as authors plagiarizing others’ work. These alternative reasons for retraction are troubling, but do not represent the same risk to downstream research integrity that retractions due to fabrication do. Because PubMed does not contain structured data about the reasons for retraction, we turned to RetractionWatch, which provides detailed structured data. This database contained 22,970 retracted papers indexed in PubMed with detailed reasons for retraction. We subset these retracted papers to those calling into question the validity of the data, such that it would be problematic if others had built upon that work. This yielded 22,955 retracted papers through November of 2025 that met our criteria as retractions that could call into question the data and results of the papers themselves. Interestingly, this suggests that retractions due exclusively to reasons other than result integrity issues are rare (18). We began by quantifying the number of retractions in biomedicine (Figure 1). The number of retracted papers has steadily increased since 2000, from a few dozen retracted papers per year to a peak of nearly 7,000 in 2023.

**Figure 1.**
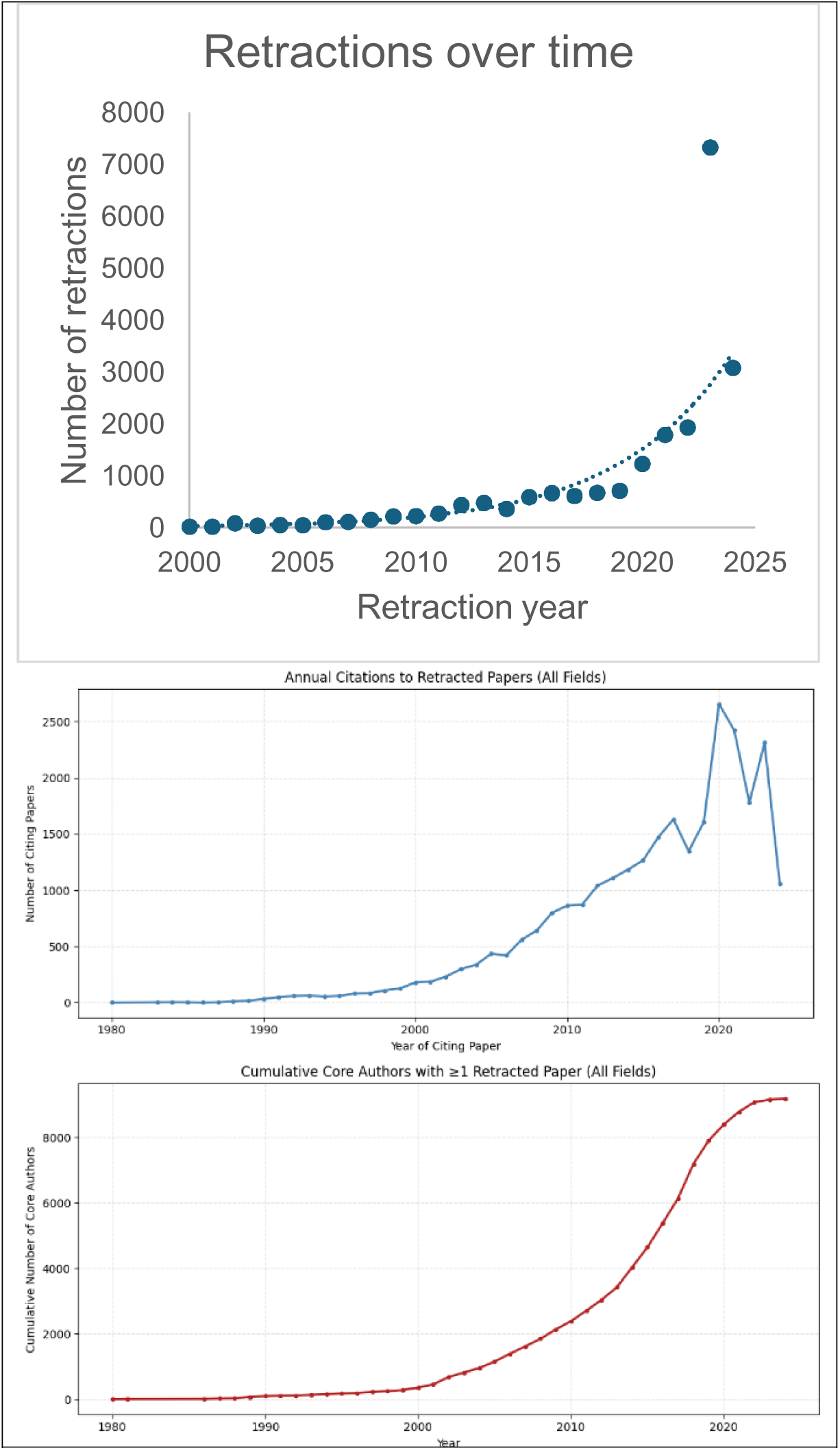
Retraction characteristics. (Top) Retractions per year due to research integrity issues, indexed in RetractionWatch and PubMed. (Middle) Increasing numbers of citations to retracted papers in recent years. (Bottom) Cumulative number of authors across the ten selected fields who have at least one retracted publication.

Our previous study examined the structural risk of retracted papers on the field of Alzheimer’s Disease research (17) to establish a framework for analyzing risk in biomedical research fields. For the present study, we expanded our analysis to ten fields of research broadly representative of different areas of biomedicine. All but one were defined using the National Library of Medicine’s Broad Subject Categories (Allergy and Immunology, Cell Biology, Drug Therapy, Genetics, Internal Medicine, Molecular Biology, Neoplasms, Neurology, and Social Sciences). We selected these fields to represent the conceptual breadth of the biomedical literature spanning from basic research to applied clinical science, and also align with areas in which the National Institutes of Health funds research. We also included COVID-19 research as a special case of a rapidly emergent field, which we identified from article Medical Subject Heading codes. This yielded a total of 5,724,015 papers, approximately 14% of the 40,000,000+ articles indexed in PubMed. Within these fields of research, 6,279 papers were in our RetractionWatch dataset (0.1% of the total, mirroring the 0.1% total for all of PubMed). Because continued use and citation of questionable knowledge from retracted papers is such an important question (5,6), we quantified the citations to retracted work in these fields. A total of 29,334 articles cited the 6,279 retracted papers across fields (Figure 1), an average of approximately five citations per retracted paper. Given the recent spike in retractions since 2020, coupled with the substantial lag in citations (19), most citations to these retracted works have probably not yet occurred.

One major issue that has been identified by previous work is the prospect of repeat retractions by individual authors. Retractions due to misconduct are thought not to be randomly distributed, but significantly cluster in individual authors (12). To address this question, we used disambiguated author profiles from the PubMed Knowledge Graph (20). This yielded 12,107,946 authors across the different fields, or an average of two per paper. The number of unique authors who have had at least one paper retracted is 26,802. Some published in multiple fields, making 32,414 author-field pairs when accounting for different field membership. Number of authors with retractions in our dataset has also risen sharply (Figure 1), a result of exponential growth of biomedicine coupled with increasing collaboration over time. If repeat retractions pe author pose a systemic risk, this base of authors constitutes a quantifiable structural risk factor. Taken together, these results point to rising retractions and potentially contaminated knowledge flow from these articles as a relatively small but increasingly important research integrity risk.

### Retraction varies across fields

Previous research indicated that biomedical research has a higher rate of retractions than other fields of research (21). Because our selected fields vary significantly in terms of their field growth, co-authorship patterns, and methodology, we asked how retraction varies across these ten selected fields. All fields except for COVID-19 showed an exponential growth trajectory. COVID-19 follows an inverted U-shaped curve, which follows the time course of the pandemic (Figure 2). The growth of the number of unique authors across these publications closely tracks the number of papers published in each field.

**Figure 2.**
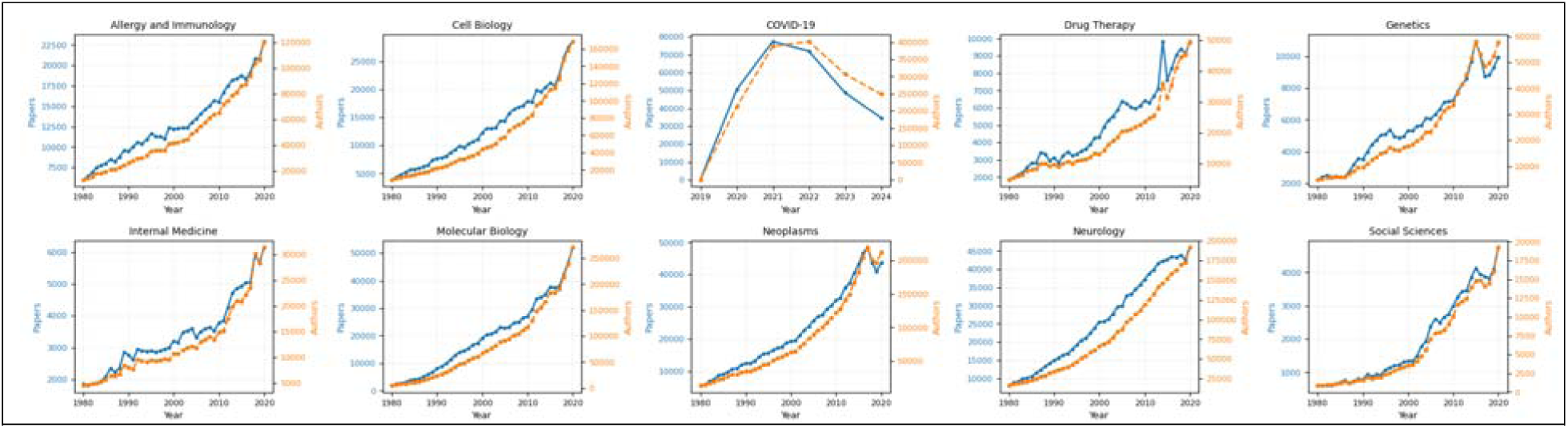
Number of papers and authors in each of the ten fields of science over time.

We next quantified the retractions by field to see how these are distributed across fields (Table 1). Retraction rates varied by over an order of magnitude, ranging from 0.02% of the literature to 0.25% of the literature. There are clear differences between fields. Cell Biology ranks first with 1,596 retracted papers and a retraction rate of 0.25%, which is also the highest among the ten fields. Neurology (1,336 papers, 0.1%), Molecular Biology (1,255 papers, 0.1%), and Neoplasm (1,177 papers, 0.11%) also have high retraction counts. These four fields together contribute over 85% of all target retracted papers in our sample, while their combined publication volume only accounts for approximately 71% of the ten-field total. By comparison, Social Sciences (16 papers, 0.02%), Genetics (89 papers, 0.03%), and Internal Medicine (77 papers, 0.05%) have significantly lower retraction counts and rates. COVID-19 constitutes a special case in that this field has produced 196 retracted papers within an observation window of only five years; however, this retraction rate of 0.076% is lower than the average for PubMed (0.1%).

**Table 1.** Retracted papers and citations to retracted papers by field.

| **Topic** | **Total Papers** | **Retracted Papers** | **Pct Retracted** | **Papers Citing Retracted** | **Pct Citing Retracted** |
| --- | --- | --- | --- | --- | --- |
| *Allergy and Immunology* | 628,289 | 291 | 0.05 | 1,368 | 0.22 |
| *COVID-19* | 258,348 | 196 | 0.08 | 2,730 | 1.06 |
| *Cell Biology* | 642,966 | 1,596 | 0.25 | 6,861 | 1.07 |
| *Drug Therapy* | 253,339 | 246 | 0.10 | 240 | 0.09 |
| *Genetics* | 274,215 | 89 | 0.03 | 142 | 0.05 |
| *Internal Medicine* | 162,659 | 77 | 0.05 | 131 | 0.08 |
| *Molecular Biology* | 1,053,520 | 1,255 | 0.12 | 5,975 | 0.57 |
| *Neoplasms* | 1,096,255 | 1,177 | 0.11 | 8,216 | 0.75 |
| *Neurology* | 1,248,446 | 1,336 | 0.11 | 3,750 | 0.30 |
| *Social Sciences* | 105,978 | 16 | 0.02 | 43 | 0.04 |

The proportion of literature citing retracted papers also varies significantly across fields. Within-field citations to retracted articles also varied by over an order of magnitude, from 0.04% (Social Sciences) to 1.1% (Cell Biology). More than 1% of the Cell Biology and COVID-19 articles cited a retracted paper. The latter effect is not necessarily because of broader disciplinary trends; the related Allergy and Immunology field as a whole only cited retracted articles 0.2% of the time. The two are nearly equal in retraction citation rate, but the mechanisms may not be identical. For COVID-19, the higher retraction citation rate is more likely related to the high-density citation network formed in a short period of time. In Cell Biology, the higher retraction citation rate is more related to its larger retraction base. By comparison, the retraction citation rates of Drug Therapy (240 papers, 0.1%), Genetics (141 papers, 0.05%), Internal Medicine (131 papers, 0.08%), and Social Sciences (43 papers, 0.04%) are all significantly lower, broadly consistent with their smaller retraction bases, lower citation density, or relatively dispersed research structure. The overall retraction rate describing the average scale of the retraction problem thus conceals the highly uneven distribution of retraction risk across fields. These results indicate that retraction risk is distributed unevenly across fields rather than occurring at random.

### Repeat author retractions pose a structural risk factor

Repeat retractions by the same author are thought to be a research integrity problem, and we next asked whether this can explain some of the field differences in retraction risk. As noted above, the ten fields contain 32,414 authors in total associated with at least one retracted paper in each field (authors unique within a field but possibly appearing in more than one field, after deduplication across fields, 26,802 authors). Cell Biology ranks first again, with a total of 8,869 retraction-associated authors; Molecular Biology (6,816) and Neoplasms (6,800) follow closely; Neurology also has 4,845 authors associated with retraction events. If retraction events are distributed approximately randomly across the author population, then historical retraction records have limited identifying value for subsequent risk. Conversely, if retraction occurs significantly more often for some authors, then retraction risk is not merely a single-paper problem but may also be related to certain authors’ sustained research practices, data-processing procedures, or experimental norms.

We therefore asked whether retractions mainly manifest as one-time, dispersed paper-quality problems, or whether they recur in the same group of authors. Our results show that multi-retraction authors (authors with two or more retracted papers) account for a non-negligible proportion of all authors of retracted papers in several fields. For example, in Neoplasms, Molecular Biology, Cell Biology, and Social Sciences the fraction of multi-retraction authors compared to all authors with at least one retraction are 11%, 9%, 9%, and 8% respectively (Table 2). In order to determine whether multiple retractions are merely a random result naturally arising from the number of retracted papers and the authorship structure, we performed a permutation test that merges the ten fields and removes cross-field duplicates. The results show that, of the 26,802 distinct authors with at least one retracted paper, 2,792 have two or more retracted papers, giving an observed proportion of 10.42%. By comparison, across 1,000 random simulations the expected multi-retraction author proportion is only 1.73%, with a 95% confidence interval of 1.52%–1.95%, corresponding to an Obs/Exp of 6.04 and a two-sided permutation p = 0.002. This result indicates that retraction events are not randomly distributed across the author network, but are significantly concentrated in individual author profiles, at a rate six times above what would be expected by chance.

**Table 2.**
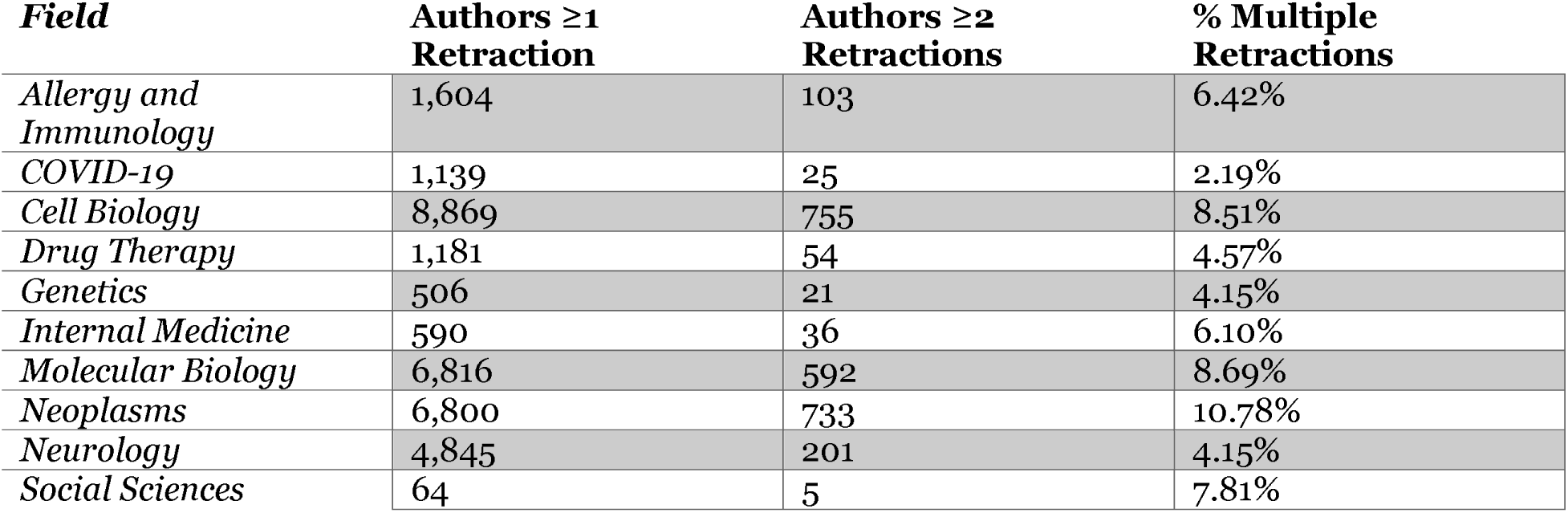
Authors with multiple retractions by field.

Previous work has suggested that newer authors are at higher risk of having work retracted than established authors (21). The roles of researchers at different career stages differ greatly, and are typically reflected in author position. First authors are typically early career, and generally conduct experiments, perform data analysis, and spearhead the progression of an individual project. Last authors are usually more established and take on responsibilities such as research design, resource coordination, team management, and quality control. When stratifying by field, we see that first-author retraction risk is higher for some, but not all fields. We calculated the percent of authors with first- vs. last-author multiple retractions (Table 3). Globally, the ratio of last:first author multiple retractions is 1.3. Social Sciences and Genetics have ratios of 1.0, indicating parity in first- vs last-author multiple retractions. Allergy and Immunology and COVID-19 have sub-1.0 last:first author multiple retraction ratios, indicating that retraction risk is lower for last authors compared to first authors. However, in four fields (Drug Therapy, Internal Medicine, Molecular Biology, and Neurology), the last:first author multiple retraction ratio was above both 1.0 and the global value of 1.3. This indicates that retraction risk stratified by author role is unevenly spread across fields of biomedical research. Together with the results of our permutation test, these findings suggest that research integrity risk may be concentrated in the portfolio of papers that are not yet retracted but have been authored by scientists with prior retractions.

**Table 3.** First- and last-authorships with multiple retractions by field.

| **Field** | **% Multiple (First Author)** | **% Multiple (Last Author)** | **Last/First Ratio** |
| --- | --- | --- | --- |
| *Allergy and Immunology* | 7.06% | 5.53% | 0.78 |
| *COVID-19* | 2.65% | 2.16% | 0.82 |
| *Cell Biology* | 6.29% | 8.07% | 1.28 |
| *Drug Therapy* | 1.33% | 4.93% | 3.7 |
| *Genetics* | 3.53% | 3.53% | 1 |
| *Internal Medicine* | 4.17% | 10.15% | 2.43 |
| *Molecular Biology* | 6.11% | 8.72% | 1.43 |
| *Neoplasms* | 6.83% | 8.48% | 1.24 |
| *Neurology* | 1.82% | 2.37% | 1.3 |
| *Social Sciences* | 8.33% | 8.33% | 1 |

We therefore further extend the vulnerability metrics to examine three categories of exposure scope: papers citing literature by authors with at least one retraction, papers citing literature by authors with multiple retractions, and papers located in clusters with retraction counts above expectation (see “Retraction risk is concentrated in specific topic clusters” section for this last category). When the risk definition is extended from “retracted papers” to “authors with retraction history,” the exposure range expands significantly. Globally, the ten fields together have approximately 1,271,550 papers with citations to papers by authors with at least one retraction, accounting for approximately 22% of the total publication volume of the ten fields. This proportion is approximately 43 times the proportion of direct citations that go to retracted papers as a fraction of all citations (0.5%). In other words, although retracted papers themselves account for only a small share of the overall literature, the entire body of work by authors with retraction history is collectively cited by a fifth of the biomedical literature.

One objection to this line of reasoning might be that this is an overly expansive definition of risk. The vast majority (90%) of authors of a retracted paper do not go on to have a second article retracted. Using a narrower risk definition (citation to authors of multiple retracted papers) reduces the scope significantly, although still at a non-negligible level. The ten fields together have approximately 333,075 papers that have cited papers by authors with multiple retraction records, accounting for approximately 6% of the total publication volume of the fields, still approximately 11 times the proportion of direct citations to retracted papers.

In our work establishing a risk analysis framework using Alzheimer’s Disease papers as a model, we examined the citation reach of individual authors in the field as a measure of risk, reasoning that the most prolific authors might pose the greatest risk to the field (17). However, it was unclear whether this mechanism of risk is plausible. Retraction risk might be lower for prolific authors rather than higher. To test this question, we next examined whether retractions occur more or less frequently than expected for prolific authors. We ranked authors within each field by publication count and separately extracted the top 1%, top 5%, and top 10% high-output author groups, and used the two-sided Fisher’s exact test to compare the number of authors with at least one retracted paper in these groups against the expected value under random allocation (Table 4). The results showed high consistency across the 30 combinations of ten fields by three percentile groups: all p-values were less than 0.005, the vast majority were less than 0.0001, and the Obs/Exp ratios were all higher than 2.0. Taking the Top 1% high-output author group as an example: the Obs/Exp for Social Sciences reaches 11.7, Neoplasms 9.2, Allergy and Immunology 8.5, Molecular Biology 8.2, Cell Biology 6.1, COVID-19 5.4, and Neurology 5.2. Even when extended to the Top 10% author group, the lowest Obs/Exp is still 2.0 (Genetics), and the highest still reaches 4.3 (Neoplasms). This shows retractions tend to occur in the most prolific author profiles regardless of field, and this finding is insensitive to the particular definition of prolific author threshold.

**Table 4.**
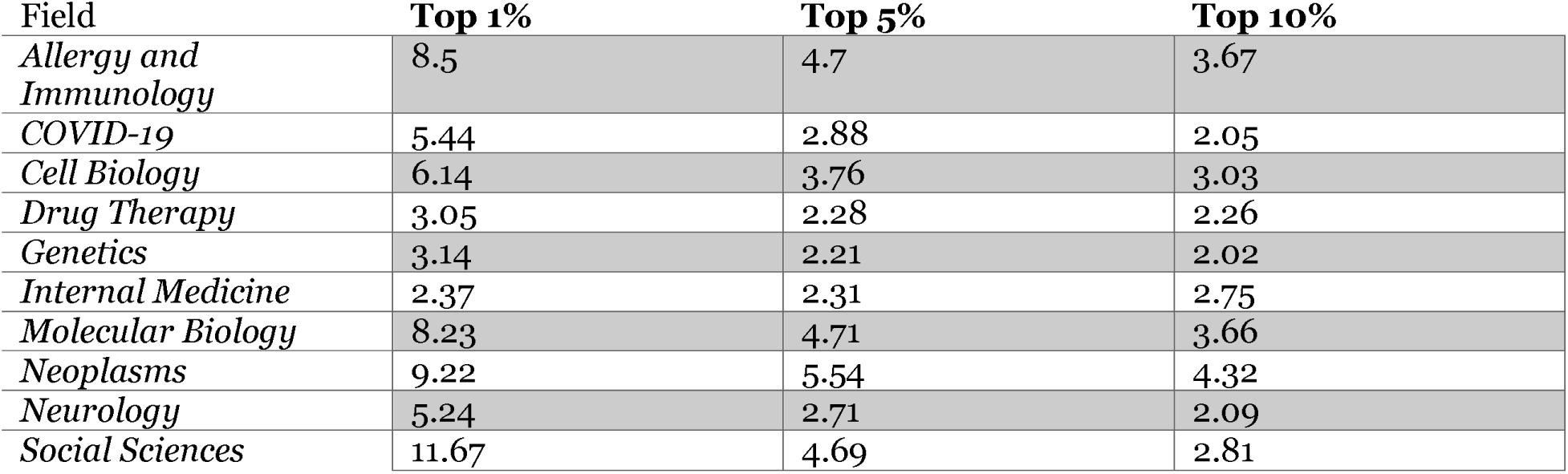
Observed retraction rates divided by expected retraction rates for most prolific authors in the top 1%, top 5%, or top 10% of number of publications in their fields.

| Field | Top 1% | Top 5% | Top 10% |
| --- | --- | --- | --- |
| *Allergy and Immunology* | 8.5 | 4.7 | 3.67 |
| *COVID-19* | 5.44 | 2.88 | 2.05 |
| *Cell Biology* | 6.14 | 3.76 | 3.03 |
| *Drug Therapy* | 3.05 | 2.28 | 2.26 |
| *Genetics* | 3.14 | 2.21 | 2.02 |
| *Internal Medicine* | 2.37 | 2.31 | 2.75 |
| *Molecular Biology* | 8.23 | 4.71 | 3.66 |
| *Neoplasms* | 9.22 | 5.54 | 4.32 |
| *Neurology* | 5.24 | 2.71 | 2.09 |
| *Social Sciences* | 11.67 | 4.69 | 2.81 |

Since retractions cluster among the most prolific authors, we next asked how much of the literature in each field cites individual authors. As in our previous study, we wanted to focus on core authors in a field, not individuals who publish only one paper (17). Likewise, because responsibility for an article falls disproportionally on the first and last authors in most fields, we focused on authors who have published at least two first- or last-author papers within a field (referred to as core authors). Measured as the proportion of within-field literature citing at least one paper by a given author, in most fields even the author with the highest coverage typically reaches only about 1%-2% of the literature (Figure 3); in fields such as Internal Medicine and Social Sciences, this proportion is even below 1%. Neoplasms is a relatively prominent exception, with its top author coverage reaching 7.5%. If the analysis scope is expanded to the top 150 high-output authors in each field, the per-author coverage further drops to only 0.1%–0.7%. Among them, both Internal Medicine and Drug Therapy are below 0.2%, while COVID-19, although the highest, does not exceed 0.7%. This suggests that our previous results indicating citation reach of 7.8% among the top 150 authors in Alzheimer’s Disease (17) were idiosyncratic to that field and represents the high end of what should be expected in biomedical research. For Neurology as a whole, the value is less than 2%.

**Figure 3.**
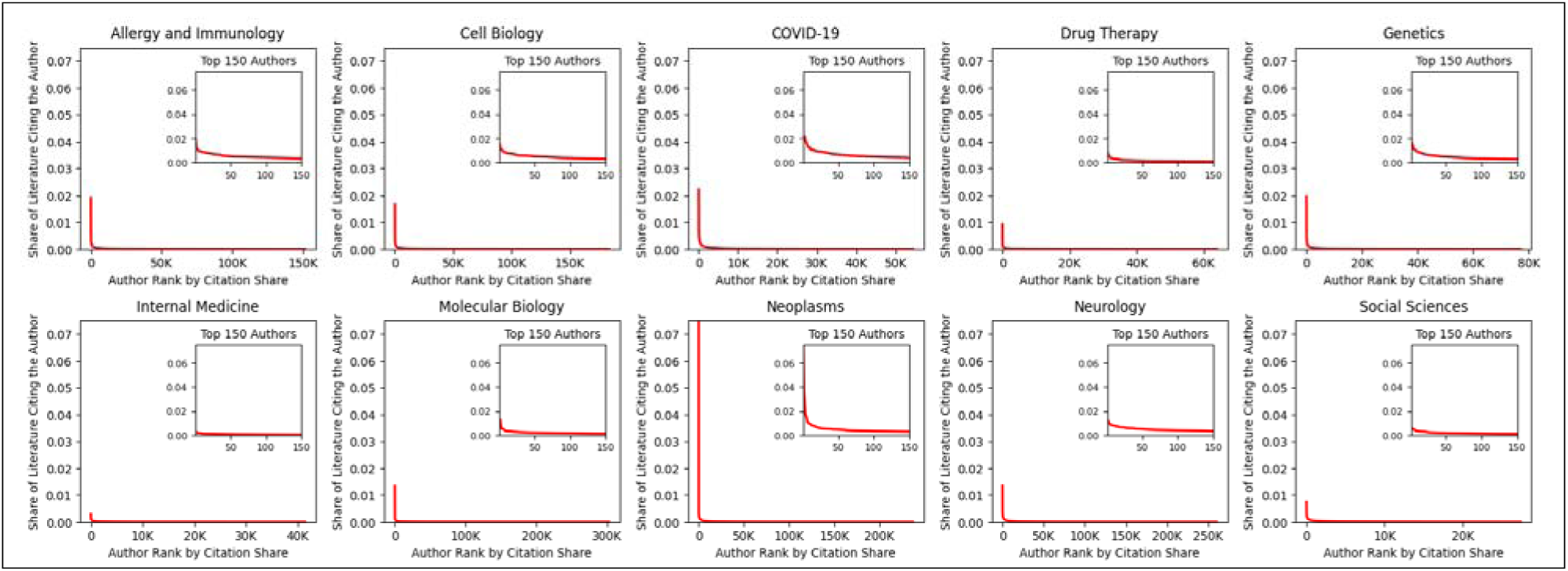
Reach of authors to the literature in ten fields of science. Main panels, fraction of field citing individual authors, sorted from the most highly cited authors in each field to the least. Inset, reach of top 150 authors in each field.

Taken together, these results demonstrate that: multiple retractions for individual authors occur much more frequently than expected by chance; the body of work particularly at risk of propagating potentially unreliable findings (citing authors with multiple retractions) is much higher than the body of work directly citing retracted papers; and retractions are particularly concentrated in the profiles of the most prolific authors.

### Retraction risk is concentrated in specific topic clusters

Our analysis thus far characterized broad field-specific differences in retraction rates, knowledge diffusion, and author-level risk quantification. The direct retraction rates of the ten fields are all below 0.25%, and papers citing retracted studies are mostly below 1%. However, this seemingly limited risk assessment has a potential bias in that the assessment scale itself may be too broad. Over the past several decades, the vast majority of life-science fields have undergone exponential expansion, and the paper volume of a single field can exceed a million papers. By contrast, the capacity of individual researchers to read, write, and cite has always been constrained. Risk may not fall evenly within field topics, just as we showed it does not fall evenly across fields. For example, Neurology as a whole has a much lower degree of single author citation than does its Alzheimer’s Disease subset. A topic-level analysis is important because it may reveal two risk factors that are difficult to identify at a field-level analysis. First, authors might acquire prominent reputations in their selected topics in a way that does not translate to an entire field. This might lead to higher within-topic citations to individual people or labs than is visible at the field level. Second, many of the theoretical causal mechanisms underlying research integrity risk have to do with subfield specific norms and practices, such as which methods to use (22), and which analytical techniques are considered appropriate (23).

In order to measure this risk at the topic level, we used the Leiden algorithm to perform clustering on the citation networks of each field (see Methods). This approach creates topically linked clusters that are connected by their citations, so topic clusters represent areas of inquiry that share common citing and referenced papers. We focused on clusters that contained at least 50 papers for subsequent analysis. The ten fields together resulted in a total of 6,863 such clusters. These contained on average 84% of the papers in each field. We began by asking how much of a topic cluster’s papers cited work from a single author. We did not select the author for each topic cluster, but instead iterated through authors and identified the most highly cited author in that topic to measure the structural vulnerability of that topic to retracted work from a single prolific author. Authors were not randomly distributed across topic clusters. Median top author citation reach ranged from 9.3% (Social Sciences) to 18.4% (Neoplasms, Table 5), much higher than top-150 author citation reach for the field as a whole (1-2%). These numbers describe the central tendency across all clusters with at least 50 papers. This is commensurate with the measurements we observed in our prior analysis of the Alzheimer’s Disease subset of the Neurology literature (17). To measure maximum vulnerability for a topic within a field, we quantified the maximum author exposure of the most vulnerable cluster within each field. This value ranged from 38.8% of a topic’s papers citing a single author (Social Sciences) to 93.9% (Allergy and Immunology). Thus, the research integrity risk seems to be highly concentrated within topics--the proportion of literature that the same author can reach within a topic cluster is more than ten times higher than the proportion they can reach in the entire field.

**Table 5.** Author reach within each field’s topic clusters. Only clusters with at least 50 papers were included. For each of these, the author with the greatest number of citations in the cluster was identified. Max author reach (maximum) reports the % of a cluster’s articles that cited the author with the most citations, but only for the cluster within each field where that value was the highest. Max author reach (median) reports the % of clusters’ articles that cited the author with the most citations for that cluster, but instead reports the median value over all clusters in a field that met the size limitation. For example, in Allergy and Immunology, the cluster with the most single-author citation exposure had 93.9% of its articles citing that author, while the median % of Allergy and Immunology clusters’ papers citing a single author was 16.7%.

| Field | Number of clusters<br>($\geq 50$<br>papers/cluster) | Max author reach<br>(maximum of all<br>clusters) | Max author reach<br>(median of all<br>clusters) |
| --- | --- | --- | --- |
| *Allergy and Immunology* | 643 | 93.90% | 16.70% |
| *COVID-19* | 254 | 59.20% | 11.90% |
| *Cell Biology* | 916 | 72.70% | 14.30% |
| *Drug Therapy* | 551 | 74.50% | 9.90% |
| *Genetics* | 393 | 81.00% | 13.30% |
| *Internal Medicine* | 335 | 42.40% | 11.90% |
| *Molecular Biology* | 1772 | 70.40% | 12.90% |
| *Neoplasms* | 842 | 68.00% | 18.40% |
| *Neurology* | 949 | 65.20% | 16.90% |
| *Social Sciences* | 208 | 38.80% | 9.30% |

What these findings reveal is the potential scope of information propagation, rather than the actual frequency at which research integrity risk has occurred. Although high author coverage indicates that these local structures have strong conditions for risk amplification, it does not mean that retracted papers have already widely permeated these clusters. To measure the extent to which these structural vulnerabilities manifest in reality, we filtered topic clusters with at least 50 papers and that had at least one retracted paper, and measured the citation reach of retracted papers in these topic clusters. Overall, median retraction citation rates were lower than the overall fraction of papers in a field citing retracted papers (less than 1% for all fields, Figure 4). However, citations to retracted papers in the most affected topic clusters was considerably higher. These ranged from 1.6% (Social Sciences) to 38% (Neurology). We therefore observe a dual effect: most topic clusters have a reduced actual share of their papers citing retracted papers, but this risk is concentrated in a small number of vulnerable clusters, which have citations to retracted articles that are an order of magnitude higher.

**Figure 4.**
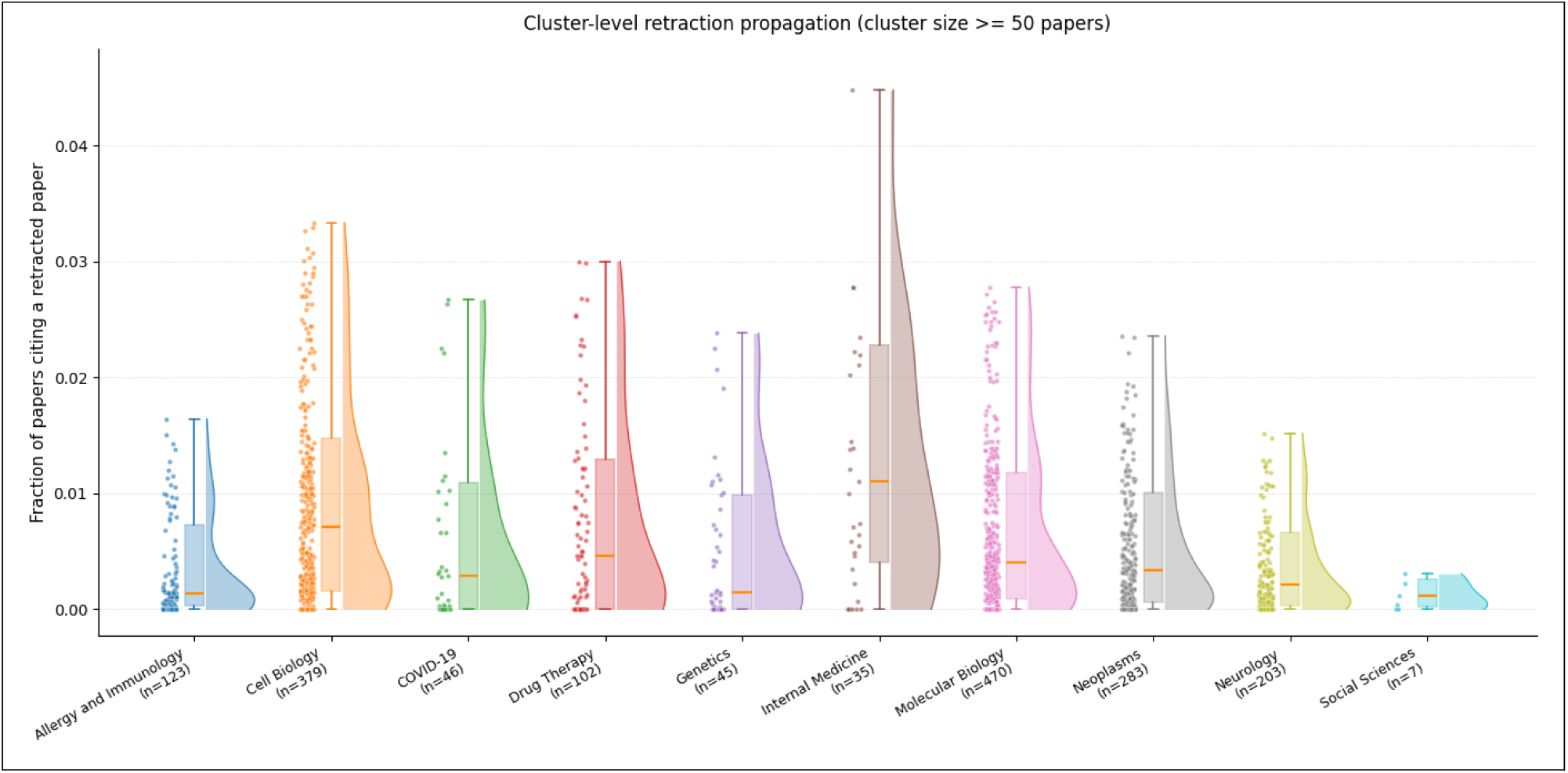
Cluster-level citations to retracted papers, for topic clusters containing at least 50 papers. Whiskers represent 1.5x the interquartile range.

To ascertain whether the risk of the underlying retraction events themselves is unevenly distributed among topic clusters, we measured the occurrence of retractions among all of the topic clusters in the ten fields. We measured the number of clusters containing at least one retracted paper, and compared this frequency with the expected number of clusters based on chance if retractions were randomly distributed (Table 6). All fields except for Social Sciences showed a lower number of clusters with retracted papers than expected by chance (0.74, Neoplasms to 0.92, Genetics, of expected numbers), indicating a concentration of retractions in some topic clusters. We therefore ran permutation tests to determine statistical significance. Social Sciences and Genetics did not have statistically significant deviations from chance, while the other fields all had significant effects, indicating significant concentration of retractions in those fields’ topic clusters.

**Table 6.** Concentration of retraction risk in each field’s topic clusters. Obs/exp ratios below 1.0 indicate that retractions are concentrating into a smaller number of clusters than expected by chance.

| *Field* | **Clusters**<br>**≥50** | **Total**<br>**Retracted** | **≥1 Obs** | **≥1 Exp** | **≥1 Exp**<br>**95% CI** | **≥1**<br>**Obs/Exp** | **≥1 p**<br>**(two-**<br>**sided)** |
| --- | --- | --- | --- | --- | --- | --- | --- |
| *Allergy and Immunology* | 643 | 232 | 123 | 141.9 | [131, 153] | 0.87 | 0.004 |
| *Cell Biology* | 916 | 1390 | 379 | 418.7 | [398, 441] | 0.91 | 0.002 |
| *COVID-19* | 254 | 146 | 46 | 61.2 | [52, 70] | 0.75 | 0.002 |
| *Drug Therapy* | 551 | 181 | 102 | 125.6 | [115, 136] | 0.81 | 0.002 |
| *Genetics* | 393 | 65 | 45 | 48.8 | [43, 55] | 0.92 | 0.3037 |
| *Internal Medicine* | 335 | 49 | 35 | 42.6 | [38, 47] | 0.82 | 0.01 |
| *Molecular Biology* | 1772 | 1149 | 470 | 579.9 | [555, 606] | 0.81 | 0.002 |
| *Neoplasms* | 842 | 1097 | 283 | 382.8 | [364, 401] | 0.74 | 0.002 |
| *Neurology* | 949 | 490 | 203 | 248 | [232, 264] | 0.82 | 0.002 |
| *Social Sciences* | 208 | 7 | 7 | 6.6 | [5, 7] | 1.06 | 1 |

Because our results indicated that multiple retractions per author occur at a rate approximately six-fold more often than expected by chance, we asked if these results were explainable by uneven distribution of multi-retraction authors, or whether topic-specific variations account for this result. If a small number of authors with multiple retracted papers are concentrated and active in specific sub-fields, they may amplify the density of retractions in these clusters. To test this possibility, within each field we removed all retracted papers under the profiles of authors with at least two retracted papers and re-performed 1,000 cluster-size-weighted permutation tests on the remaining retracted papers (Table 7). The results show that of the 8 originally statistically significant fields, after removing multi-retraction authors, the cluster-level deviation is no longer significant for Cell Biology and Internal Medicine. This indicates that the cluster concentration observed in those fields is largely driven by the cross-topic repeated publishing of multi-retraction authors. However, in six fields, Allergy and Immunology (0.89, p = 0.014), COVID-19 (0.79, p = 0.018), Drug Therapy (0.86, p = 0.002), Molecular Biology (0.91, p = 0.002), Neoplasms (0.80, p = 0.002) and Neurology (0.89, p = 0.004), after removing multi-retraction authors, the deviation remains significant. This indicates that most fields have local structural effects independent of authors’ repeat retractions. These analyses reveal that the citation network structure is itself a focal point of retraction risk. Topic clusters concentrate retracted work beyond that expected by chance, creating a vulnerability that persists even after accounting for unevenly distributed multi-retraction authors.

**Table 7.**
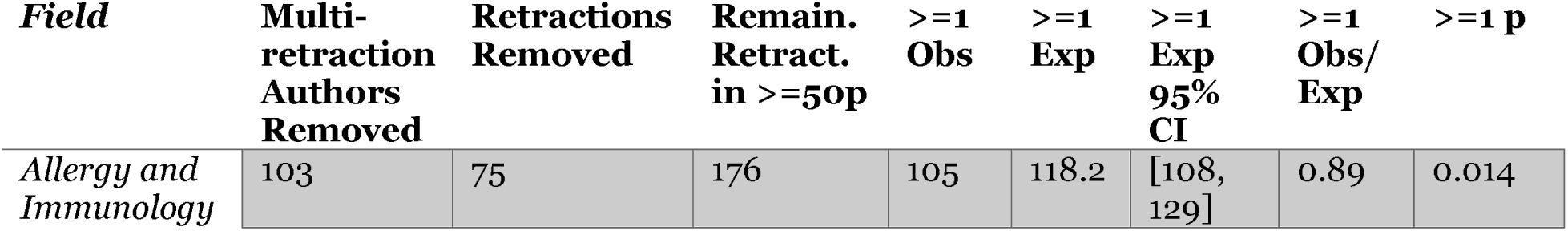
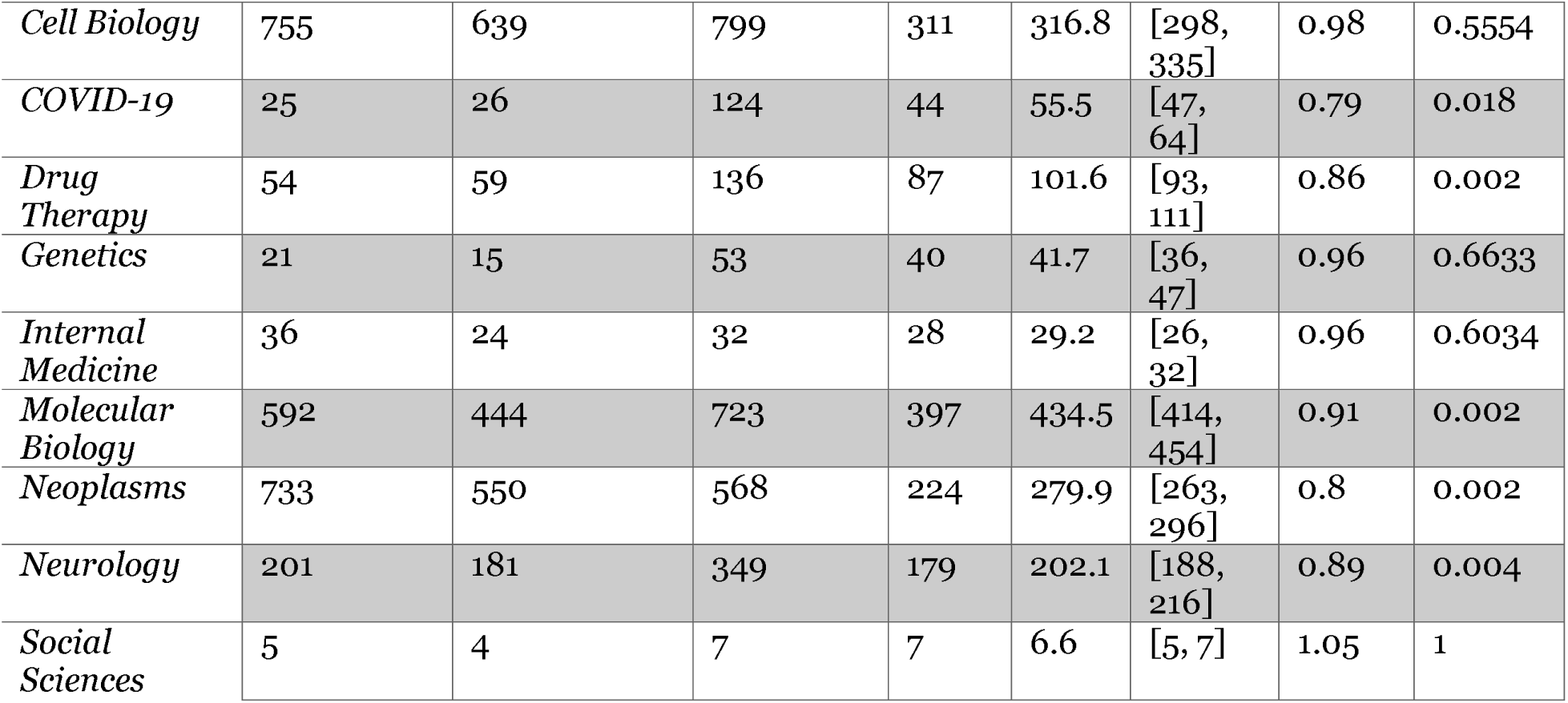
Concentration of retraction risk in each field’s topic clusters after removing the effect of authors with multiple retractions. Obs/exp ratios below 1.0 indicate that retractions are concentrating into a smaller number of clusters than expected by chance.

## Discussion

Taken together, these results demonstrate four kinds of structural research integrity risk. First, multiple retractions from individual authors occur much more frequently than would be expected by chance. Second, the citation structure of research topics within fields leads to much higher author level citation reach than field-level analysis might indicate. Third, there is a positive association between retraction risk and author productivity. Combined with the high author-level citation reach within topics, this leads to a concentration of risk within specific topics that is realized by a small number of topics containing a disproportionate number of citations to retracted papers. Finally, topic-specific risk factors are detectable, and lead to concentration of retracted papers in specific topics that is both higher than chance and largely independent of multi-retraction authors. Much of the literature has focused on structural vulnerabilities attributable to incentive structures (14), in particular a publish-or-perish environment (15) reflecting hypercompetition in biomedicine (15,24). After retraction, papers continue for years to accrue citations from new articles by authors seemingly unaware of the retracted status of the cited article (5,6). These lay bare a fundamental concern about retracted papers: they pose a research integrity risk not primarily because their individual findings are deemed questionable, but because the body of work that built upon them is suddenly uncertain. If the magnitude of this research integrity risk is structural rather than incidental, it means that it cannot be simply dismissed because retractions are, at present, rare events.

Here, our data give both cause for concern as well as grounds for cautious reassurance. In all fields examined, a small number of prolific authors reach a large share of the literature within their topics. In addition, the positive relationship between author productivity and retraction rate indicates that the conditions for amplification of unreliable findings are in place. A striking 22% of the literature cites authors who have at least one paper retracted, and 6% cites authors with multiple retractions. However, the average topic cluster cites retracted work no more frequently than the field baseline, so this amplification risk seems contained to a small number of topics. This suggests that the latent risk inherent in this organization of the scientific literature, while real and quantifiable, has not yet materialized.

The Reducing the Inadvertent Spread of Retracted Science team developed rigorous recommendations regarding the handling of retractions, including consistent information dissemination, standardization of retraction taxonomies, and stakeholder education about retractions (25). Given our findings, there are further implications for focusing efforts on minimizing research integrity risks. One is already in place by the scientific community: increased sensitivity of multi-retraction authors’ work. Robust coverage of retractions from prolific authors seems to be the norm in academia already (3,4,26–28). Retractions of highly cited articles are newsworthy to the scientific community, and this norm should be preserved. A second is to establish risk indicators of scientific topics. These are not the same for every field, according to our analysis. This might be because some fields have different standards on reporting guidelines, for example, or use different methods (23) that carry uneven levels of risk. A third would be to retrospectively analyze propagation risk for fields with high degrees of author citation reach compared to others, as in the Alzheimer’s Disease topic (17). Finally, the scientific community would particularly benefit from targeted efforts at testing and enhancing reproducibility in high-risk scientific topics.

This work is not without limitations. Most importantly, the interpretation of the multi-retraction authors should not be overstated. Retractions for work that the originating lab subsequently identifies to be unreliable can be interpreted as a positive for research integrity, rather than a negative. This critical reassessment is an important component of the self-correcting nature of science. It should be lauded, not punished. A second limitation is that citation reach of authors should not be conflated with contamination of the literature, even for studies citing a retracted paper. Not all citations indicate a substantive contribution of a cited article on a new project (29,30). Most citations are rhetorical in nature, helping to establish background information for readers or position a new discovery in relation to its field. These would not be a primary vector for calling later projects into question. Third, there may be a small number coding errors in the RetractionWatch dataset, since human curation is not perfect, and in addition, no single source of retracted article lists has been established as fully comprehensive. Finally, other analytical design choices here such as the PubMed scoping, iCite dataset or choice of clustering decisions may influence our results.

Because our dataset contains many newly retracted papers, the full extent of citations to these articles will not be visible for years to come (5,10,19). In addition, papers in our corpus that have not been retracted will be in the future. This limits our view into the full scope of the research integrity risk in the literature. However, it also provides an opportunity to improve the resilience of the system before potential risks are fully realized.

## Methods

### Data Sources and Processing

Author information was drawn from the PubMed Knowledge Graph 2.0 (20). This dataset integrates multiple types of biomedical scientific entities, including papers, patents, and clinical trials, and provides author-level information such as unique author identifiers, authorship order, and the total number of authors per paper.

Retraction information was obtained from the Retraction Watch database (31). We retained only retraction records with valid PubMed identifiers, and matched the set of retracted PMIDs against the literature sets of each research domain to identify retracted papers within each domain.

Article-level citation impact indicators, including Relative Citation Ratio (RCR) and raw citation counts, were obtained from the iCite database (32). The iCite database hosts the NIH Open Citation Collection (OCC) and article-level metrics, including RCR scores, APT (Approximate Potential to Translate) scores, and citation counts (19,33–35). RCR normalizes citation impact by comparing a paper’s citation rate against the median citation rate of NIH-funded papers published in the same year and field, where a value of 1.0 represents the median impact level within that field (Hutchins et al., 2016). Compared to journal-level metrics, article-level metrics more fairly reflect the actual academic impact of most researchers (36). Citation relationship data were drawn from the NIH hereafter OCC, accessed via iCite database snapshots (32). The OCC uses PMIDs as identifiers and provides open citation relationships among biomedical publications.

Based on OCC data, we extracted directed citation relationships within each research field, defined by the National Library of Medicine’s Broad Subject Categories and constructed domain-level citation networks for subsequent analyses.

For the COVID-19 field, OCC data also effectively cover citation relationships between preprints and peer-reviewed articles, although since there is no Broad Subject Category for COVID-19, these papers were identified with Medical Subject Header terms as we previously used for identifying Alzheimer’s Disease research (17).

### Author Filtering

After generating domain-level aggregate statistics, this study further focused the analysis on authors who had published at least two papers as first author or last author over the course of their careers. In the ten domains covered by this study, according to general academic conventions in the biomedical field, the first author typically leads experimental design, data analysis, and manuscript preparation, while the last author typically takes responsibility for research supervision, resource coordination, and project organization; both authorship positions can therefore be regarded as reflecting substantive leadership contribution to the research. In the PubMed Knowledge Graph (20), author number zero is a placeholder value, and was removed from analysis.

The author filtering procedure was as follows: first, we retained paper–author pairs in which the PMID belonged to the target domain’s literature set and the author unique identifier was nonzero; we then retained only records in which the author appeared as either first or last author; finally, applying a threshold of “having published at least two papers in either of these two roles,” we aggregated qualifying authors into an eligible author set. This eligible author set was used for the author-level citation-reach analyses (i.e., the fraction-citing and citation-reach results in Figure 3). The retraction-related author analyses—retraction concentration, conditional retraction probability, the Fisher’s exact test for prolific authors, and the global multi-retraction author baseline—were instead computed over all authors with a nonzero unique identifier whose papers fall within the domain’s literature set, without restricting to first/last-author roles or applying the ≥2-paper threshold. For these analyses, ‘prolific’ authors were defined by total publication count across all authorship positions (top 1%/5%/10%). It should be noted that these filtering criteria apply only to author-level statistical analyses; citation network construction and clustering analyses are still based on all papers in each domain’s literature set and are not subject to the author filtering criteria.

The rationale for this filtering criterion is that authors who have published only one paper in a given domain are often difficult to classify as genuinely sustained contributors to that field, as opposed to having only a one-time or marginal association with it over the course of their careers. By contrast, authors who have published at least two papers as first or last author more clearly demonstrate sustained research engagement and stable academic participation in the field. Therefore, to improve the domain representativeness and result stability of the author-level analyses, only authors meeting the above criteria were retained for subsequent analyses.

Following filtering, the number of authors included in the analysis for each domain was as follows: Allergy and Immunology 151,320; Cell Biology 183,459; COVID-19 54,438; Drug Therapy 63,867; Genetics 77,096; Internal Medicine 41,386; Molecular Biology 302,722; Neoplasms 236,861; Neurology 260,467; and Social Sciences 27,415.

### Citation Network Construction

#### Citation Edge Preprocessing

For the directed citation edges in each domain, we first removed records in which the citing and cited papers were the same article, to exclude self-citation effects. Each directed edge was then converted into an undirected canonical form sorted by PMID magnitude, and duplicate edges were removed, yielding an undirected citation graph for each domain to serve as input for subsequent clustering analyses.

### Citation Clustering Analysis

#### Clustering Method

Following citation edge preprocessing for each domain, this study employed the Leiden community detection algorithm to perform thematic clustering of the literature (37), rather than semantic clustering (38,39). Citation network clustering, as a general methodological framework for identifying breakthrough advances in biomedical research, has been successfully applied to the analysis of knowledge structures in large-scale biomedical literature (17,40,41). To avoid generating clusters that are either too fragmented or too coarse, and to achieve a balance between cluster interpretability and structural stability, the resolution parameter was uniformly set to 0.0003. Clustering results were subsequently mapped back to individual papers via PMID, to support subsequent retraction propagation analysis and author coverage analysis.

#### Cluster Eligibility Filtering

In retraction propagation analysis, if a retracted paper appears in a cluster containing very few papers, its citation diffusion ratio can be artificially inflated to near 100%, thereby obscuring true propagation patterns. To reduce the interference of micro-clusters, this study used at least 50 papers within the cluster as the criterion for eligible (large) clusters; subsequent related indicators were computed independently within large clusters (≥50 papers) and small clusters (<50 papers).

#### Retraction Distribution Analysis at the Cluster Level

This study matched the PMIDs of retracted papers in the Retraction Watch dataset against the cleaned paper PMIDs in each domain, identified retracted papers within each domain, computed retraction propagation indicators at the cluster level, and tested whether retracted papers exhibit clustering in specific knowledge communities beyond what would be expected by chance. This analytical framework also draws on the field vulnerability assessment method proposed in (17), which quantifies the potential negative impact of fraudulent researchers on an entire research field by examining the extent to which their citation networks have penetrated the field’s literature.

To determine whether the distribution of retracted papers across large clusters exhibits clustering tendencies beyond random expectation, within each domain we randomly redistributed the retracted papers across all eligible clusters with ≥50 papers 1,000 times, weighted by cluster size (multinomial distribution), and computed the expected values under random conditions for both number of clusters containing ≥1 retracted paper and number of clusters containing ≥2 retracted papers. We then compared these expected values to the observed values and computed Obs/Exp ratios; we also computed two-sided p-values (taking twice the smaller of the proportions of permutation values in each extreme tail) and 95% confidence intervals (2.5th–97.5th percentiles of the permutation distribution) to assess the statistical significance of the deviation of observed values from random expectation. An Obs/Exp value <1 indicates that retracted papers are more concentrated across large clusters than expected by chance, while a value >1 indicates greater dispersion. All random procedures used a fixed random seed of 42 to ensure reproducibility.

#### Conditional Test for Multi-Retraction Authors

To further assess whether the cluster-level retraction distribution observed is driven solely by multi-retraction authors, this study employed a conditional test: we first identified multi-retraction authors (those with two or more retracted papers) within each domain and removed all of their retracted papers from the cluster-level retraction counts; we then reran 1,000 cluster-size-weighted permutation tests on the remaining non-multi-retraction-author retracted papers, computing Obs/Exp, two-sided p-values, and 95% confidence intervals as described above. If the cluster-level deviation remained significant after removing multi-retraction authors, this suggests the existence of local structural effects in that domain that are independent of authors’ repeated retraction behavior.

#### Identification of Clusters with Above-Expected Retractions

As a complement to the vulnerability indicators, this study identified large clusters with above-expected retraction counts. Within each domain, the expected value was defined as [total retractions in large clusters / total number of large clusters] (ceiling), and clusters with retraction counts exceeding this expected value were flagged as above-expected clusters. The total number of papers within these clusters and their share of all papers in large clusters were computed to measure the range of papers covered by more vulnerable local structures.

#### Author-Level Retraction Analysis

To assess whether retractions are concentrated among particular author groups, this study first identified all authors within each domain associated with at least one retracted paper based on their AID, and computed the proportion of such authors with two or more retracted papers, as a measure of the degree of concentration of retraction risk at the individual level. To examine whether there are systematic differences in retraction patterns across author roles, we separately computed the conditional probability of a subsequent retraction for first authors and corresponding authors who had already experienced at least one retraction.

#### Prolific Authors and Retraction Concentration

To test whether retractions are concentrated among prolific authors, we first ranked all authors within each domain in descending order of publication count, and extracted the top 1%, top 5%, and top 10% of prolific authors. For each quantile group, we counted the number of authors with at least one retracted paper and compared this to the expected value under random assignment (expected value = total retraction-associated authors in the domain x corresponding quantile proportion). The statistical test used was a two-sided Fisher’s exact test. The Obs/Exp ratio indicates the magnitude of actual retraction concentration relative to random expectation, and the p-value assesses the statistical significance of this concentration. All tests were conducted independently across the 30 combinations formed by ten domains and three quantile groups. Prior research has shown that in NIH-funded scientific domains, research topic choice is systematically associated with funding outcomes (38); this study similarly tests whether publication volume and retraction risk exhibit a quantifiable structural pattern.

#### Global Multi-Retraction Author Baseline Estimation

To assess the overall tendency toward repeated retraction at a broader scale, this study estimated the proportion of multi-retraction authors and their random baseline at the global level by pooling all ten domains. We identified all authors in the union of the ten domains associated with at least one retracted paper (AID nonzero) and computed the proportion with two or more retracted papers as the observed value. A random baseline was simultaneously constructed by randomly drawing without replacement, from all author–paper pairs (restricted to AID nonzero and PMID belonging to any of the target domains), a number of papers equal to the global retraction count to serve as a simulated retraction set, then recomputing the proportion of multi-retraction authors. This process was repeated 1,000 times to obtain the expected value and 95% confidence interval under random conditions, and a two-sided permutation test was used to assess the statistical significance of the deviation of the actual proportion from random expectation.

## Notes

### Competing Interest Statement

The authors have declared no competing interest.

## References

1. Cokol M, Ozbay F, Rodriguez-Esteban R. Retraction rates are on the rise. EMBO Rep. 2008 Jan;9(1):2. doi:10.1038/sj.embor.7401143 PubMed PMID: 18174889; PubMed Central PMCID: PMC2246630.

2. Else H. Biomedical paper retractions have quadrupled in 20 years — why? Nature. 2024 Jun 13;630(8016):280–1. doi:10.1038/d41586-024-01609-0

3. Piller C. Researchers plan to retract landmark Alzheimer’s paper containing doctored images. Science. 2024 Jun 4. doi:10.1126/science.zdo1zpb

4. Piller C. Blots on a field? Science. 2022 Jul 22;377(6604):358–63. doi:10.1126/science.add9993 PubMed PMID: 35862524.

5. Schneider J, Ye D, Hill AM, Whitehorn AS. Continued post-retraction citation of a fraudulent clinical trial report, 11 years after it was retracted for falsifying data. Scientometrics. 2020 Dec;125(3):2877–913. doi:10.1007/s11192-020-03631-1

6. Hsiao TK, Schneider J. Continued use of retracted papers: Temporal trends in citations and (lack of) awareness of retractions shown in citation contexts in biomedicine. Quant Sci Stud. 2022 Feb;2(4):1144–69. doi:10.1162/qss_a_00155 PubMed PMID: 36186715; PubMed Central PMCID: PMC9520488.

7. Bolland MJ, Grey A, Avenell A. Citation of retracted publications: A challenging problem. Account Res. 2022 Jan;29(1):18–25. doi:10.1080/08989621.2021.1886933 PubMed PMID: 33557605.

8. Bar-Ilan J, Halevi G. Post retraction citations in context: a case study. Scientometrics. 2017;113(1):547–65. doi:10.1007/s11192-017-2242-0 PubMed PMID: 29056790; PubMed Central PMCID: PMC5629243.

9. Lou Y, Zhou Z, Li M. Beyond Retractions: A Topic-Based Analysis of the Impact and Lifespan of Retracted Papers. Journal of Data and Information Science. 2026 Jun 26. doi:10.1515/jdis-2026-0064

10. Hwang SY, Yon DK, Lee SW, Kim MS, Kim JY, Smith L, et al. Causes for Retraction in the Biomedical Literature: A Systematic Review of Studies of Retraction Notices. J Korean Med Sci. 2023 Oct 23;38(41):e333. doi:10.3346/jkms.2023.38.e333 PubMed PMID: 37873630; PubMed Central PMCID: PMC10593599.

11. Casadevall A, Steen RG, Fang FC. Sources of error in the retracted scientific literature. FASEB J. 2014 Sep;28(9):3847–55. doi:10.1096/fj.14-256735 PubMed PMID: 24928194; PubMed Central PMCID: PMC5395722.

12. Steen RG, Casadevall A, Fang FC. Why Has the Number of Scientific Retractions Increased? Derrick GE, editor. PLoS ONE. 2013 Jul 8;8(7):e68397. doi:10.1371/journal.pone.0068397

13. Franzen M, Rödder S, Weingart P. Fraud: causes and culprits as perceived by science and the media. Institutional changes, rather than individual motivations, encourage misconduct. EMBO Rep. 2007 Jan;8(1):3–7. doi:10.1038/sj.embor.7400884 PubMed PMID: 17203094; PubMed Central PMCID: PMC1796756.

14. Fanelli D. How many scientists fabricate and falsify research? A systematic review and meta-analysis of survey data. PLoS One. 2009 May 29;4(5):e5738. doi:10.1371/journal.pone.0005738 PubMed PMID: 19478950; PubMed Central PMCID: PMC2685008.

15. Edwards MA, Roy S. Academic Research in the 21st Century: Maintaining Scientific Integrity in a Climate of Perverse Incentives and Hypercompetition. Environ Eng Sci. 2017 Jan 1;34(1):51–61. doi:10.1089/ees.2016.0223 PubMed PMID: 28115824; PubMed Central PMCID: PMC5206685.

16. Giray L, Fabros B, Xavierine J. Scientific retractions: causes, processes, and implications for research integrity. Naunyn Schmiedebergs Arch Pharmacol. 2026 May 1. doi:10.1007/s00210-026-05410-w PubMed PMID: 42065765.

17. Ni C, Hutchins BI. Framework for assessing the risk to a field from fraudulent researchers: A case study of Alzheimer’s disease. Asso for Info Science & Tech. 2025 Sep;76(9):1162–73. doi:10.1002/asi.25009

18. Fang FC, Steen RG, Casadevall A. Misconduct accounts for the majority of retracted scientific publications. Proc Natl Acad Sci U S A. 2012 Oct 16;109(42):17028–33. doi:10.1073/pnas.1212247109 PubMed PMID: 23027971; PubMed Central PMCID: PMC3479492.

19. Hutchins BI, Yuan X, Anderson JM, Santangelo GM. Relative Citation Ratio (RCR): A New Metric That Uses Citation Rates to Measure Influence at the Article Level. PLoS Biol. 2016 Sep;14(9):e1002541. doi:10.1371/journal.pbio.1002541 PubMed Central PMCID: PMC5012559.

20. Xu J, Yu C, Xu J, Torvik VI, Kang J, Sung M, et al. PubMed knowledge graph 2.0: Connecting papers, patents, and clinical trials in biomedical science. Sci Data. 2025 Jun 17;12(1):1018. doi:10.1038/s41597-025-05343-8

21. Ioannidis JPA, Pezzullo AM, Cristiano A, Boccia S, Baas J. Linking citation and retraction data reveals the demographics of scientific retractions among highly cited authors. Bandrowski A, editor. PLoS Biol. 2025 Jan 30;23(1):e3002999. doi:10.1371/journal.pbio.3002999

22. Weiskirchen R. Misidentified cell lines: failures of peer review, varying journal responses to misidentification inquiries, and strategies for safeguarding biomedical research. Res Integr Peer Rev. 2025 Jul 11;10(1):12. doi:10.1186/s41073-025-00170-2 PubMed PMID: 40640915; PubMed Central PMCID: PMC12247328.

23. von Bubnoff A. Experimental Quality [Internet]. Burroughs Wellcome Fund; 2020. Available from: https://www.bwfund.org/wp-content/uploads/2020/06/BWF_CDG_EQ_5.2.16_0.pdf

24. Lynn Kamerlin SC. Hypercompetition in biomedical research evaluation and its impact on young scientist careers. International Microbiology. 2015;(18):253–65. doi:10.2436/20.1501.01.257

25. Schneider J, Woods ND, Proescholdt R, RISRS Team. Reducing the Inadvertent Spread of Retracted Science: recommendations from the RISRS report. Res Integr Peer Rev. 2022 Sep 19;7(1):6. doi:10.1186/s41073-022-00125-x PubMed PMID: 36123607; PubMed Central PMCID: PMC9483880.

26. Rong LQ, Leshem E, Kaplan JA. Further Retractions of Dr. Joachim Boldt in the Journal of Cardiothoracic and Vascular Anesthesia. Journal of Cardiothoracic and Vascular Anesthesia. 2025 Feb;39(2):360–3. doi:10.1053/j.jvca.2024.11.016

27. Dyer O. Departing Stanford president retracts two Science papers after investigation. BMJ. 2023 Sep 4;382:2035. doi:10.1136/bmj.p2035 PubMed PMID: 37666519.

28. Enserink M. ‘We are embarrassed’: Scientific rigor proponents retract paper on benefits of scientific rigor. Science. 2024 Sep 25. doi:10.1126/science.zkz6doq

29. Teplitskiy M, Duede E, Menietti M, Lakhani KR. How status of research papers affects the way they are read and cited. Research Policy. 2022 May;51(4):104484. doi:10.1016/j.respol.2022.104484

30. Hoppe TA, Arabi S, Hutchins BI. Predicting substantive biomedical citations without full text. Proc Natl Acad Sci USA. 2023 Jul 25;120(30):e2213697120. doi:10.1073/pnas.2213697120

31. Retraction Watch Data [Internet]. https://gitlab.com/crossref/retraction-watch-data: GitLab; 2025. Available from: https://gitlab.com/crossref/retraction-watch-data

32. iCite, Hutchins BI, Santangelo GM. iCite Database Snapshots (NIH Open Citation Collection). Figshare; 2019.

33. Hutchins BI. A tipping point for open citation data. Quantitative Science Studies. 2021;1–5. doi:10.1162/qss_c_00138

34. Hutchins BI, Baker KL, Davis MT, Diwersy MA, Haque E, Harriman RM, et al. The NIH Open Citation Collection: A public access, broad coverage resource. PLoS Biol. 2019 Oct;17(10):e3000385. doi:10.1371/journal.pbio.3000385 PubMed Central PMCID: PMC6786512.

35. Hutchins BI, Davis MT, Meseroll RA, Santangelo GM. Predicting translational progress in biomedical research. PLoS Biol. 2019 Oct;17(10):e3000416. doi:10.1371/journal.pbio.3000416 PubMed Central PMCID: PMC6786525.

36. Arabi S, Ni C, Hutchins BI. Most researchers would receive more recognition if assessed by article-level metrics than by journal-level metrics. PLoS Biol. 2025 Dec;23(12):e3003532. doi:10.1371/journal.pbio.3003532 PubMed PMID: 41401134; PubMed Central PMCID: PMC12707641.

37. Traag VA, Waltman L, Van Eck NJ. From Louvain to Leiden: guaranteeing well-connected communities. Sci Rep. 2019 Mar 26;9(1):5233. doi:10.1038/s41598-019-41695-z

38. Hoppe TA, Litovitz A, Willis KA, Meseroll RA, Perkins MJ, Hutchins BI, et al. Topic choice contributes to the lower rate of NIH awards to African-American/black scientists. Sci Adv. 2019 Oct;5(10):eaaw7238. doi:10.1126/sciadv.aaw7238 PubMed Central PMCID: PMC6785250.

39. Afshar AS, Yang Q, Thebault-Spieker J, Hutchins BI. Quantifying (mis)alignment between reader focus and editor citation in scholarly biomedical topics in Wikipedia. Quantitative Science Studies. 2026 Mar 13;1–23. doi:10.1162/QSS.a.457

40. Arabi S, Hutchins BI. Forecasting novel therapeutic development in biomedical research. bioRxiv. 2026 Jun 1. doi:10.64898/2026.05.29.728775

41. Davis MT, Busse BL, Arabi S, Meyer P, Hoppe TA, Meseroll RA, et al. Prediction of transformative breakthroughs in biomedical research. bioRxiv. 2025 Dec 17;2025.12.16.694385. doi:10.64898/2025.12.16.694385 PubMed PMID: 41624054; PubMed Central PMCID: PMC12857589.

